# A genome-wide RNA interference screening reveals protectiveness of SNX5 knockdown in a Parkinson’s disease cell model

**DOI:** 10.1101/2024.03.13.584847

**Authors:** Matthias Höllerhage, Linghan Duan, Oscar Wing Ho Chua, Claudia Moebius, Svenja Bothe, Franziska Hopfner, Christian Wahl-Schott, Marc Bickle, Günter U. Höglinger

## Abstract

**Background:** Alpha-synuclein is a major player in the pathophysiology of a group of diseases called synucleinopathies, which include Parkinson’s disease, dementia with Lewy bodies, and multiple system atrophy. To date, there is no disease-modifying therapy available for these synucleinopathies. Furthermore, the intracellular mechanisms by which alpha-synuclein confers toxicity are not yet fully understood. Therefore, it is of utmost importance to investigate the pathophysiology of alpha-synuclein-induced toxicity in order to identify novel molecular targets for the development of disease-modifying therapies.

**Methods:** In the present study, we performed the first genome-wide siRNA modifier screening in a human postmitotic neuronal cell model using alpha-synuclein-induced toxicity as read-out. In a multi-step approach, we identified several genes, whose knockdown protected from alpha-synuclein-induced toxicity. The main hit was further validated by different methods, including immunofluorescence microscopy, qPCR, and Western blot.

**Results:** The highest protection was achieved by knockdown of *SNX5*, which encodes the SNX5 protein, a component of the retromer complex. We confirmed the protective efficacy of *SNX5* knockdown with an independent siRNA system. SNX5 protein is part of SNX-BAR heterodimers, which are part of the retromer complex. We found that extracellular and overexpressed intracellular alpha-synuclein led to fragmentation of the trans-Golgi network, which was prevented by *SNX5* knockdown by confining alpha-synuclein in early endosomes.

**Conclusion:** In summary, our data suggest that SNX5 plays an important role in trafficking and toxicity of alpha-synuclein. Therefore, SNX5 appears to be a possible target for therapeutic interventions in synucleinopathies.

## Background

Synucleinopathies are a group of neurodegenerative diseases, defined by the presence of intracellular proteinaceous inclusions consisting mainly of aggregated alpha-synuclein (αSyn). In case of PD and dementia with Lewy bodies (DLB), these αSyn aggregates are found in neurons and designated as Lewy bodies (1). Multiple system atrophy is characterized by αSyn aggregates, called glial cytoplasmic inclusions (GCI), in oligodendrocytes (2). Physiologically, αSyn is small, unfolded and soluble protein with 140 amino acids. However, in synucleinopathies, αSyn aggregates and confers neuronal toxicity. In PD, the typical motor symptoms (bradykinesia, rigidity, and tremor) are caused by the death of dopaminergic neurons in the substantia nigra pars compacta. It is widely accepted that αSyn plays a role in synapses (3). The mechanisms, how αSyn becomes toxic are not fully understood. However, a common assumption suggests that small oligomeric species occurring in the aggregation process are toxic for neuronal cells (3). Furthermore, the release of αSyn into the extracellular space and the uptake of αSyn species into neighboring cells are believed to play a role in cell-to-cell spreading of αSyn pathology throughout the brain as a mechanism involved in progression of the disease (4). In order to investigate αSyn pathophysiology, we established a model in differentiated, postmitotic dopaminergic Lund human mesecenphalic (LUHMES) neurons (5), in which the moderate overexpression of human wild-type αSyn leads to ∼50% cell death upon, which is accompanied by the occurrence of a 37 kDa oligomeric species of αSyn that was not present in control cells (6). In this model, we have previously investigated intracellular mechanism involved in αSyn degradation and identified protective interventions (7–9). In the present work, we performed a genome-wide siRNA screening in this model to identify genes, whose knockdown protects against αSyn-induced toxicity. The most interesting hits was sorting nexin 5 (SNX5), a member of the retromer complex, which is a cargo recognition complex involved in endosome sorting and endosome-trans Golgi network (TGN) trafficking (10).

Since mutations in vacuolar protein sorting ortholog 35 (VPS35), another component of the retromer complex, are associated with hereditary forms of PD (11) and endocytosis and endosomal trafficking have been linked to uptake and release of αSyn (12), our focus was to investigate the involvement of SNX5 in regulation and trafficking of αSyn and how SNX5 gene knockdown protected dopaminergic neurons from αSyn-induced cytotoxicity.

### Aim

The aim of the study was to perform a genome-wide siRNA screening in order to identify novel molecular targets for a potentially disease-modifying therapy of neurodegenerative synucleinopathies. Furthermore, we aimed to perform a deeper characterization of the main hit from this screening.

## Methods

### Cell culture

In all experiments, LUHMES cells (5) were cultured at 37 °C, 5% CO_2_, and 100% humidity, as previously described (8, 9). Briefly, for proliferation, LUHMES cell were cultured in growth medium (GM) consistent of DMEM/F12 (Sigma-Aldrich, St. Louis, MO, USA), 1% N2 supplement (Life Technologies, Carlsbad, CA, USA), and 0.04µg/ml basic fibroblast growth factor (bFGF, PeproTech, Rocky Hill, CT, USA) in cell culture flasks (Nunc, Thermo Fisher Scientific, Waltham, MA, USA). Before seeding of the cells, the flask pre-coated with poly-L-ornithine (0.1 ml/ml, incubation overnight at 37 °C) and washed three times with phosphate buffered saline (PBS; Life Technologies, Carlsbad, CA, USA). For differentiation, the cells were seeded in differentiation medium (DM) consistent of DMEM/F12, 1% N2 supplement, 1-µg/ml tetracycline (Sigma-Aldrich), 0.49-µg/ml dibutyryl cyclic adenosine monophosphate (Sigma-Aldrich), and 2 ng/ml glial cell-derived neurotrophic factor (GDNF, R&D Systems, Minneapolis, MN, USA). The experiments were conducted on flasks or multi-well plates (Nunc) double-coated with PLO as described above followed by a second coating with fibronectin (5µg/ml, incubation overnight at 37 °C; R&D Systems, Minneapolis, MN, USA) and once washing with PBS.

### Adenoviral transduction

Adenoviral vectors of serotype 5 (BioFocus, Charles River Laboratories Nederland B.V., Leiden, Netherlands) were used to overexpress human wild-type αSyn or GFP under a cytomegalovirus promotor. In the high-content screening, in which the cells were seeded in GM, a MOI of 5 was used to account for proliferation of the cells on the day after seeding. Twenty-four hours after transduction remaining viral vectors were removed by three times washing with PBS. In the experiments to investigate the pathophysiology, the cells were transduced with the vectors two days after seeding in DM with a multiplicity of infection (MOI) of 2.

### Genome-wide RNA interference screening

In the high-throughput screening, the cells were prepared as we did in a previous compound screening (9). Briefly, cells were seeded in double-coated flasks in GM with a medium change to DM after 24 hours. One day after the initiation of differentiation, the cells were transduced with adenoviral vectors to overexpress human αSyn. One day thereafter remaining virus particles were removed by rinsing the cells with PBS and then the cells were detached using trypsin-EDTA (5 min incubation at 37 °C; Sigma-Aldrich) and reseeded in double-coated 384-well multi-well-plates one day after transduction. By doing so, we achieved maximal homogeneity throughout the screening plates. After seeding in the multi-well plates the cells were treated with the different endoribonuclease-prepared small interfering RNAs (esiRNAs). esiRNAs are a RNAi system with high efficiency and small off-target effects (13, 14). Cell death was quantified as the percentage of cells with propidium iodide (PI) incorporation, cell survival as cells without PI incorporation. PI is an intercalating compound that is actively removed from living cells and therefore accumulates only in dead cells. For the quantification, the cells were incubation with 4 μg/ml PI (PI, Sigma-Aldrich) and 2 μg/ml Hoechst 33342 to stain all cell nuclei. Imaging was performed after 15 min with an Opera High Content Screening System (Perkin Elmer, Waltham, MA, USA). In the whole screening, a genome-wide library containing esiRNAs against 16,744 genes was tested. Cells transfected with an esiRNA against αSyn were used as positive controls (best survival), cells transfected with an esiRNA against firefly luciferase (F-Luc), and mock transfected cells were used as negative control. In the primary screening, the survival of the cells after transfection with all individual esiRNAs was compared to the average survival of the whole library and Z-scores were calculated (Z-statistics). Primary hits were defined as esiRNAs that led to a survival Z-score > 2.4 in at least two screening runs (2x Z > 2.4) compared to the whole library. In total 46,627 individual experiments were performed. In the secondary screening, the cells were seeded, transduced with adenoviral vectors, transfected with the esiRNA, and imaged in the same way. In this screening, three runs were performed and an ANOVA followed by a Dunnett’s post-hoc test was used to compare the survival rate. All esiRNAs that led to a significantly better survival than the F-Luc esiRNA (p-value < 0.05) were considered as secondary hits. In the tertiary screening, the survival of the cells transfected with these esiRNAs was investigated again in GFP-expressing cells and the survival was compared to the survival in αSyn-overexpressing cells. Those esiRNAs that specifically protected against αSyn as determined by multiple T-tests were considered as final hits.

### siRNA transfection

The cells were transfected with either MISSION^®^ esiRNA (Sigma-Aldrich) at a concentration of 200 ng/µl or siPOOL siRNA (SiTOOLs Biotech, Munich, Germany) at a concentration of 5 nM. Before transfection the siRNAs were mixed with OptiMEM medium (Thermo) and Lipofectamine RNAiMax (2 µl/ml; Thermo Fisher Scientific) and incubated at RT for 20 min.

### Quantitative real-time polymerase chain reactions

For quantitative real-time polymerase chain reactions (qPCR), RNA was collected by using the RNeasy Minit kit (Qiagen, Venlo, Netherlands) and the RNA concentration was determined with a NanoDrop 2000 (Thermo Fisher Scientific) spectrometer. Subsequently, the RNA samples were reverse transcribed into cDNA using the iScript cDNA Synthesis kit (Bio-Rad Laboratories). The qPCR was performed with a CFX96 Touch Real-Time PCR Detection System (Bio-Rad Laboratories), using SYBR Select (Thermo Fisher) as dye with. The following primers were used in this study: SNX1: forward: AAGCACTCTCAGAATGGCTTC, reverse: CGGCCCTCCGTTTTTCAAG; SNX2: forward: GGGAAGCCCACCGACTTTG, reverse: GGCCATTGGAGTTTGCACTAATA; SNX5: forward: TCTGTATCTGTGGACCTGAATGT, reverse: GTGGGCAGTGTGGTCTTTGT); SNX6: forward: TCTTTGAGCACGAACGAACA, reverse: CATCAGCAGCACTTTTGTGAG; VPS35: forward: GTCAAGTCATTTCCTCAGTCCAG, reverse: CCCCTCAAGGGATGTTGCAC.

### Western blot

For Western blot, whole cell extracts were collected by using M-PER lysis buffer (Thermo Fisher Scientific) supplemented with protease inhibitor and phosphatase inhibitor cocktail (Merck Millipore, Burlington, MA, USA) on ice. After centrifugation (15,000 g for 10 min at 4 °C), the supernatants were collected and protein content was measured using the BCA Protein assay (Thermo Fisher Scientific) according to the manufacturer’s instructions. 20 µg protein in Laemmli Sample Buffer (Bio-Rad Laboratories, Hercules, CA, USA) per lane were loaded on 4-12% bis-tris gels (Bio-Rad Laboratories) or 4-15% Tris-glycine gels (Bio-Rad). After sodium dodecyl sulfate polyacrylamide gel electrophoresis (SDS-PAGE), the proteins were blotted onto methanol-activated polyvinylidene difluoride (PVDF) membranes (0.2 µm) with a Trans-Blot SD Semi-Dry Transfer Cell system (Bio-Rad Laboratories). After blotting, the membranes were fixed by incubation with paraformaldehyde (PFA; 0.4%) for 30 min. After rinsing with tris-buffered saline with 0.05% Tween-20 (TBS-T, pH 7.4), the membranes were blocked with 3x Rotiblock for 1h at RT. Thereafter, the membranes were incubated with the primary antibodies in 1x Rotiblock in TBS-T overnight at 4 °C, rinsed three times with TBS-T followed by incubation with a corresponding horse radish peroxidase (HRP)-conjugated secondary antibody in 1x Rotiblock in TBS-T for 2h at RT. After three times rinsing with TBS-T, the membranes were incubated with Clarity ECL substrate (Bio-Rad) for 10-15 min at RT. Images were taken with a LI-COR Odyssey® Fc imaging system (LI-COR Biotechnology, Lincoln, NE). Loading control was performed by incubation of the membranes with a β-actin or a glyceraldehyde 3-phosphate dehydrogenase (GAPDH) antibody. The following primary antibodies were used in this study: mouse anti-SNX1 (1:1000; Santa Cruz Biotechnology; Dallas, TX, USA), mouse anti-SNX2 (1:1000; Santa Cruz Biotechnology), mouse anti-SNX5 (1:1000; Santa Cruz Biotechnology), mouse anti-SNX6 (1:1000; Santa Cruz Biotechnology), goat anti-VPS35 (1:1000; Abcam), rabbit anti β-actin (Cell Signaling Technology), rabbit anti-GAPDH (1:2000; Cell Signaling Technology). The following secondary antibodies were used: HRP-coupled anti-goat, -mouse, or -rabbit antibody (1:5000; Vector Laboratories, Burlingame, CA, USA)

### Immunocytochemistry

Sterile glass coverslips (Bellco Glass, Vineland, NJ, USA) in 24-well plates (Nunc) or ibidi dishes (ibidi; Gräfelfing, Germany) were double-coated as described above. For immunocytochemistry (ICC), the cells were fixed by incubation with paraformaldehyde (PFA; 4%) for 30 min at RT followed by a permeabilization step by using Triton X-100 (0.1%; Sigma-Aldrich) for 15 min at RT or Tween-20 (0.1%) for 5 min at RT. After permeabilization, the cells were washed three times with PBS and then blocked with normal horse serum (NHS; 5%; Vector Laboratories) for 1h at RT or overnight at 4 °C. Thereafter, the cells were incubated with the primary antibodies in NHS (5%) overnight at 4 °C, washed three times with PBS followed by incubation with the secondary antibodies for 2h at RT. Ten minutes before the end of the incubation period, 4′,6-diamidino-2-phenylindole (DAPI; 1 μg/ml final concentration) was added to the cells. After another three times washing with PBS, images were obtained. The following primary antibodies were used in this study: anti-tubulin III (1:000; Santa Cruz Biotechnology), rabbit anti-TGN46 (1:1000; Abcam, Cambride, UK), mouse anti-Rab5a (1:100; Santa Cruz Biotechnology; Dallas, TX, USA), rabbit anti-Rab7 (1:100; Abcam, Cambride, UK), rabbit anti-LAMP1 (1:1000; Abcam, Cambride, UK), rabbit anti-LAMP2a (1:200; Abcam, Cambride, UK), rabbit anti-p62 (1:50; Abcam, Cambride, UK), rabbit anti-LC3B (1:400; Abcam, Cambride, UK), rabbit anti-Rab11a (1:1000; Invitrogen). The following secondary antibodies were used: Alexa Fluor 488 donkey anti-mouse, or anti-rabbit antibody (1:1000; Thermo Fisher Scientific), Alexa Fluor 594 donkey anti-mouse, or anti-rabbit antibody (1:1000; Thermo Fisher Scientific).

### Lactate dehydrogenase release measurement

Six days after transduction, the cell culture medium was collected to quantify lactate dehydrogenase (LDH) that was released into the cell culture medium as a measure for cytotoxicity. Briefly, 30 µl of conditioned medium were mixed with a reaction buffer consistent of 74.24 mM Tris/HCl, 185.6 mM NaCl, 3.2 mM pyruvate, and 4 mM nicotinamide adenine dinucleotide (NADH) in water. To measure the LDH release, the NADH turnover was quantified repeated by measurements of the absorbance at 340 nm over 4 min with a FLUOstar Omega (BMG Labtech, Ortenberg, Germany).

### Treatment with labeled αSyn

Recombinant monomeric αSyn (18 mg/ml) was incubated with ATTO-565-N-hydroxysuccinimidyl-ester (ATTO-TEC, Siegen Germany) in a sodium bicarbonate buffer according to the manufacturer’s instructions to fluorescently label αSyn. After the incubation, the concentration of the resultant solution containing labelled αSyn was adjusted to 2 mg/mL and excess unbound dye was removed using Bio-Spin 6 size exclusion spin columns (Bio-Rad Laboratories). ATT565-αSyn was added to LUHMES cells on day six in vitro at a final concentration of 2 µM. After 24h, αSyn was removed by washing the cells three times with PBS. To remove labeled αSyn on the outside of the cells, the cells were incubated with trypsin-EDTA for 30 s at 37 °C followed by another three times washing with PBS.

### Microscopy and image analysis

All microcopy imaging was performed with a Leica DMi8 inverted fluorescence microscope (Leica Camera AG, Wetzlar, Germany) and Leica Application Suite (LAS) X as software. The density of the neuronal network was determined in images stained with the anti-tubulin III antibody by measurement of the total branch length as measure for network density and the number of quadruple points as measure for network complexity with a modified version of the ‘neurite analyzer’ plugin (15) in Fiji (Fiji Is Just ImageJ) version 2.14.0 for Windows 64-bit (https://imagej.net/software/fiji/). To quantify the amount of αSyn inside and outside the TGN, a line was draw across the TGN or Golgi region in each image using LAS X, and the software automatically generated an intensity profile. In each cell analyzed, four regions of interest (ROIs) lines were selected to acquire intensity profiles from both the trans-Golgi network (TGN) and adjacent areas. Quantitative analysis was conducted by measuring the area under the curve of the signal intensities inside and outside the TGN region using Fiji. A minimum of 50 cells were evaluated for each experimental condition.

To analyze the colocalization, regions of interest (ROI) were selected by drawing a line across the TGN or Golgi region and the degree of colocalization between αSyn and TGN was quantified using the Fiji software version 2.14.0 for Windows 64-bit with the JACop plugin, which calculated the thresholded Manderson’s overlap coefficient (M1t), with thresholds set automatically by the plugin (16).

The different states of the TGN (normal, scattered, and fragmented) were microscopically determined as previously described (17). A normal TGN has an intact ring shape. A scattered TGN shows a less intensely stained ring shape with some fragments separated from the ring. A fragmented TGN does not show a ring shape, but also fragments visible in the cytoplasm.

### Statistical analysis

Statistical analysis was performed using GraphPad Prism 10.0 (GraphPad Software, La Jolla, CA, USA). All datasets were tested for normality using the D’Agostino and Pearson omnibus normality test. Unless otherwise specified, a one-way ANOVA analysis of variance was performed, followed by a Tukey’s post-hoc test or pairwise comparisons between selected groups. P-values below 0.05 were considered as statistically significant.

## Results

### Genome-wide esiRNA screening identified *SNX5* knockdown as being protective against αSyn induced toxicity

In a PD cell model, in which postmitotic dopaminergic LUHMES neurons degenerate upon moderate overexpression of human wild-type αSyn (8, 6, 9), we performed a genome-wide esiRNA screening with step-wise validation. In the primary screening, esiRNAs against 16,774 genes were tested for their protective efficacy against αSyn-induced toxicity in duplicates or triplicates, leading to 46,627 experiments in total. Primary hits being selected by a Z-statistic threshold. A secondary screening was conducted to confirm the protectiveness of the hits from the primary screening vs. a non-specific control-siRNA. A third screening identified those esiRNAs protecting significantly against αSyn-induced toxicity compared to GFP-expressing cells to exclude non-specific effects on cell viability. A flowchart of the screening process is illustrated in **Fig. 1a**.

**Figure 1:**
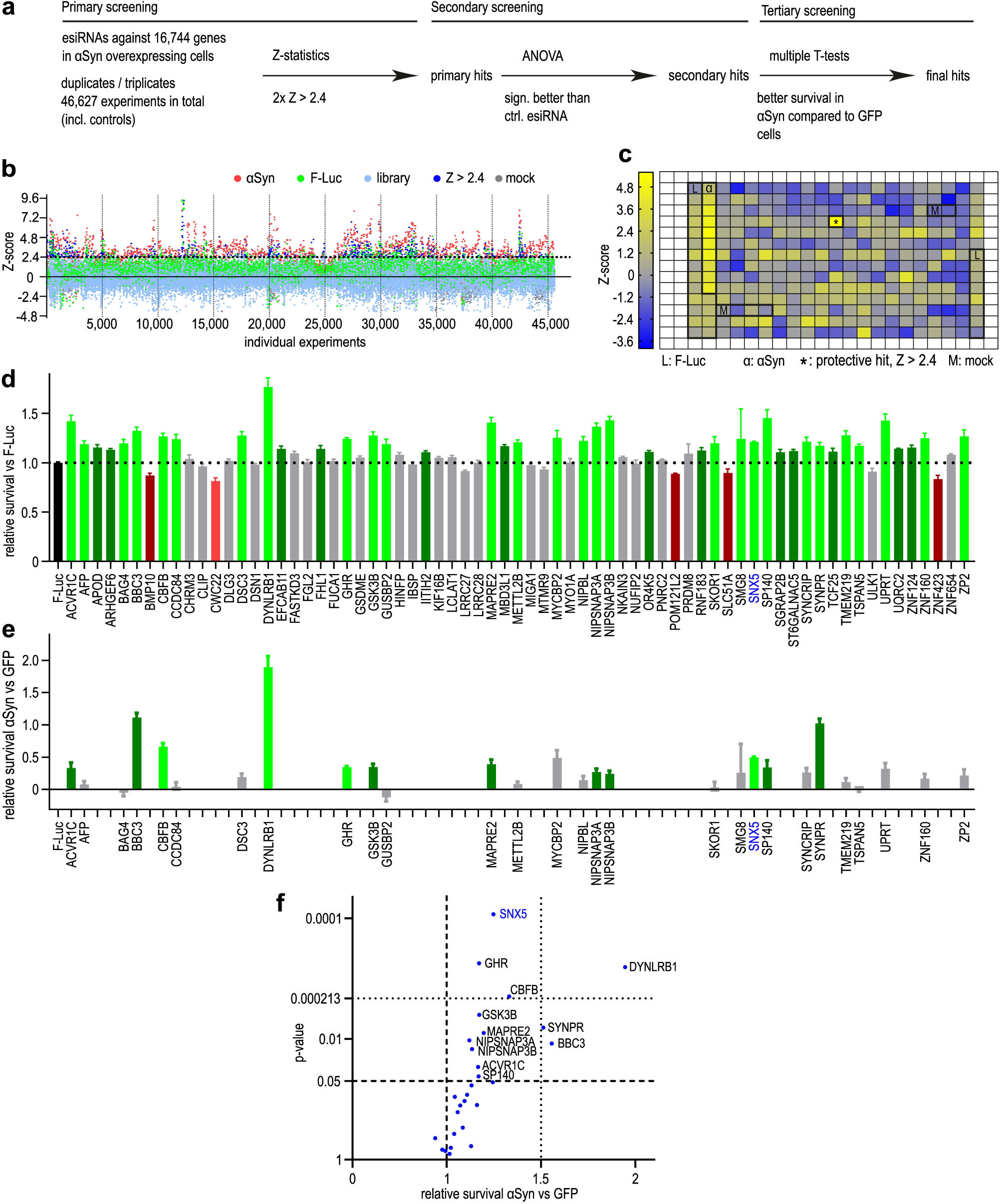
Genome-wide siRNA screening. **a:** Flowchart of the screening process. Primary screening: Z-statistics (Z > 2.4 vs. library) to identify primary hits, defined as Z > 2.4 in at least 2 screening runs (primary hits: 2x Z > 2.4). Secondary screening: ANOVA analyses (vs. F-Luc as control esiRNA). Tertiary screening, final hits: multiple T-tests (comparison to survival in GFP overexpressing cells). **b**: Z-scores of all experiments from the primary screening. Red dots: Z-scores of cells transfected with esiRNA against αSyn as positive control; green dots: Z-scores of cells transfected with esiRNA against luciferase (F-Luc) as negative control; light blue dots: Z-scores from the library, dark blue dots: Z-score > 2.4; grey dots: mock transfection. **c**: Representative results from one screening plate. Thicker lines mark areas with controls (L: luciferase, M: mock transfection, α: αSyn transfection). * indicates a protective hit (Z > 2.4). **d**: Secondary screening results: Relative survival compared to the survival of controls (esiRNA against F-Luc). Green bars: all genes the knockdown of which led to increased survival (without correction for multiple testing); light green bars: hits after correction for multiple testing (Dunnett’s post hoc test); red bars: all genes the knockdown of which led to reduced survival; light red bars: genes the knockdown of which led to significant reduction of survival after correction for multiple testing; grey bars: genes the knockdown of which had no significant influence on the survival. **e**: Comparison of the survival between αSyn-overexpressing cells and GFP-expressing cells. Green bars: all genes the knockdown of which led to a higher survival in αSyn-overexpressing cells; light green bars: significantly higher survival in αSyn overexpressing cells compared to GFP-expressing cells, after correction for multiple testing. **f**: Volcano plot showing the relative survival between αSyn-overexpressing and GFP-expressing cells. Lower dotted line: p-value < 0.05 in the individual t-tests, the upper dotted line: p-value adjusted for multiple testing. SNX5 showed the smallest p-value of all screened genes.

Positive hits were defined as esiRNAs leading to a better survival of αSyn overexpressing cells with a z-Score > 2.4 above the mean survival. In the entire screening, in 1,580 wells an esiRNA against αSyn was tested (positive control). Of these, 1,246 led to a survival with a Z-score > 2.4 (true positive rate: 79%) and 334 did not meet hit criteria (false-negative rate: 21%). The average Z-score of survival of cells transfected with the esiRNA against αSyn was 3.3 ± 1.8 (standard deviation; SD). An esiRNA against firefly luciferase (F-Luc) was used as negative control. In total, 3,964 wells were transfected with the F-Luc esiRNA, 3,566 of which yielded a Z-score < 2.4 (true negative rate: 90%), whereas 398 showed a Z-score > 2.4 (false positive rate: 10%). The average Z-score of cells transfected with a F-Luc esiRNA was 1.2 ± 1.2, suggesting a minor unspecific and non-significant protective effect of the transfection procedure itself. In 1,851 wells, a mock transfection was performed, yielding Z-scores < 2.4 in 1,849 wells (true negative rate 100%) with an average Z-score of −0.4 ± 1.0 (SD), showing no effect on cell survival. The results from the complete screening are illustrated in **Fig. 1b**. The results of an exemplary single plate are displayed in **Fig. 1c**.

In the primary screening, 80 of the 16,744 esiRNAs led to a higher cell survival in αSyn-overexpressing cells and were therefore selected per our definition as primary hits. The secondary screening was performed with 69 esiRNAs, since 11 of the 80 primary hits could not be matched to a protein, were matched to a pseudogene, or were matched to a non-coding RNA. After correction for multiply testing, 28 esiRNAs led to a significant higher protection compared to the F-Luc esiRNA control. (**Fig. 1d**). These secondary hits were tested again in αSyn- vs. GFP-overexpressing cells. Transfection with 12 thereof protected specifically from αSyn-induced toxicity (T-tests), and 4 thereof (esiRNA against *SNX5*, *DYNLRB1*, *CEFB*, and *GHR*) remained significant after correction for multiple testing (**Fig. 1e,f)**.

The esiRNA with the lowest p-value targeted *SNX5*. Therefore, we decided to further elucidate the role of *SNX5* in our PD cell model.

### Confirmation of the protective efficacy of *SNX5* knockdown against αSyn-induced toxicity using siPOOL siRNAs

After the screening process, we aimed to confirm the protective efficacy of *SNX5* knockdown against αSyn-induced toxicity again with siPOOL siRNAs. Therefore, LUHMES cells were transduced to overexpress αSyn and transfected with *SNX5*-siRNAs. The knockdown effect and the protective efficacy were determined four and six days after transduction, respectively (**Fig. 2a**). The siPOOL siRNAs efficiently reduced *SNX5* mRNA levels by 93.1 ± 42.2% (p<0.001) and also decreased SNX5 protein levels by 46.7 ± 6.1% (p<0.001) compared to a negative control siRNA (**Fig. 2b,2c**). As one readout to quantify cell viability, we performed neuronal network analyses (**Fig. 2d**). In untransduced cells, the total branch length was 37.2 ± 1.8 mm. Upon αSyn overexpressing, this was reduced to 30.2 ± 1.7 mm (p<0.05). However, knockdown of *SNX5* led to a protection of the neuronal network with total branch lengths of 40.4 ± 0.5 mm (p<0.001 vs. untransfected and control siRNA transfected cells; **Fig 2e**). In a similar manner the number of quadruple points as measure for network complexity was reduced upon αSyn overexpression (77.6 ± 7.7 vs. 107.6 ± 9.2 in untransduced control cells; p<0.05). Furthermore, *SNX5* knockdown led to conservation of the network complexity (121.3 ± 3.2 quadruple points), which was not observed with a negative control siRNA (Fig. 2f). Additionally, we quantified LDH release into the cell culture medium as measure for cell death. In αSyn-overexpressing cells (100%), the knockdown of *SNX5* reduced the LDH released into the cell culture medium by 22.4 ± 6.1% (p<0.001), which was not observed after transfection with a negative control siRNA (**Fig. 2g**).

**Figure 2:**
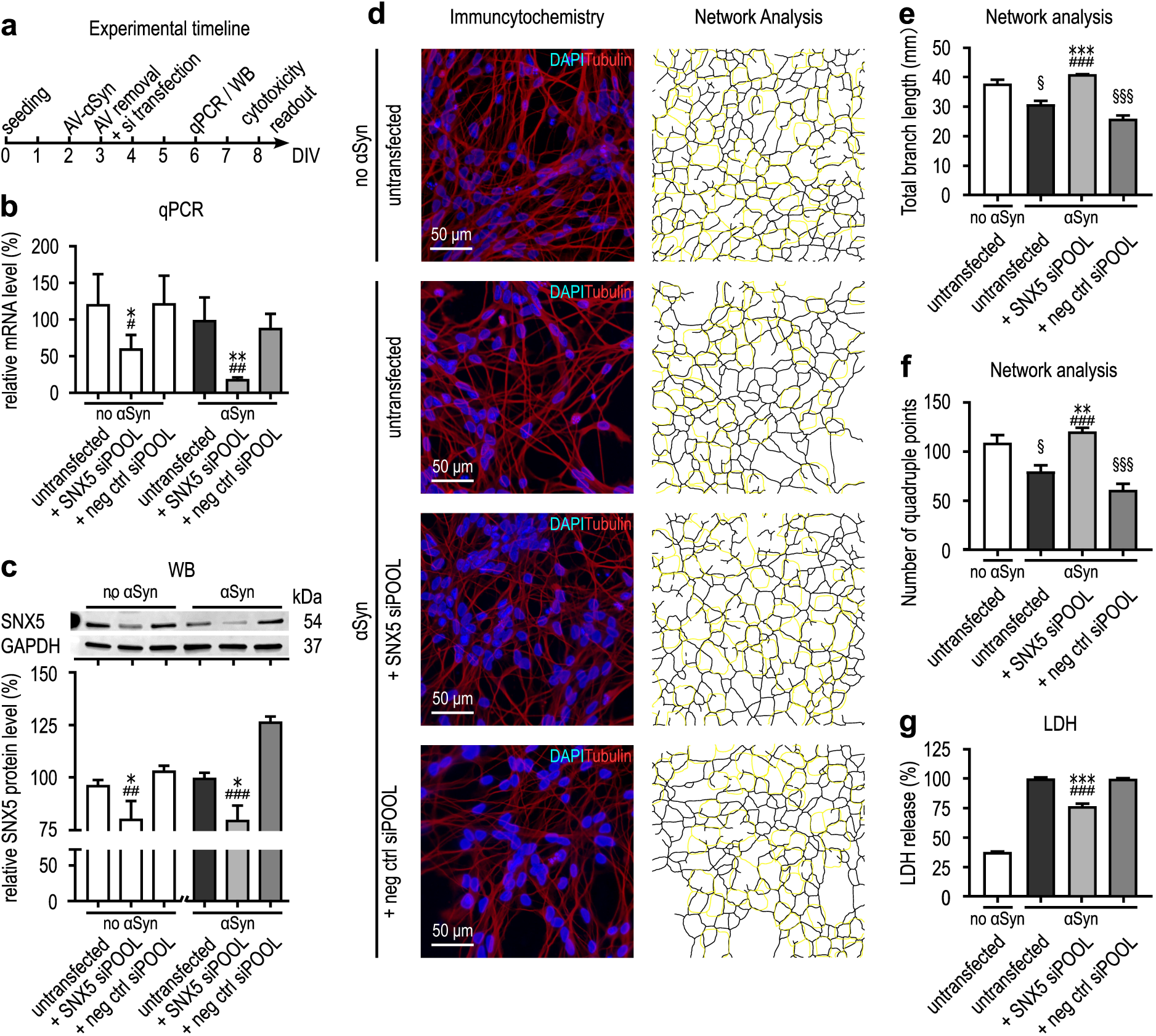
Validation of the protective efficacy of SNX5 knockdown. **a**: Experimental timeline of the validation experiment. AV: adenovirus, DIV: days in vitro **b**: Quantification of *SNX5* mRNA levels in control cells without αSyn overexpression (white bars) and in αSyn overexpressing cells (black/grey bars) in untransfected cells, cells transfected with siPOOL siRNA against SNX5, or cells transfected with negative control (neg ctrl) siPOOL siRNAs. qPCR: quantitative polymerase chain reaction. **c**: Upper panel: Representative Western blot (WB) with an antibody against SNX5 of cell without αSyn overexpression and αSyn-overexpressing cells, without siRNA transfection (untransfected), after transfection with an siPOOL siRNA against *SNX5* and after transfection with a negative control (neg ctrl) siRNA. Lower panel: quantification of the Western blots, representatively shown above. * *p* < 0.05, ** *p* < 0.01, *** *p* < 0.001 vs untransfected cells, ^#^ *p* < 0.05 ^##^ *p* < 0.01, ^###^ *p* < 0.001 vs cells transfected with a negative control siRNA. **d**: panels on the left: immunofluorescence images after staining of the neuronal network with an antibody against tubulin III (red) and nuclear co-staining with DAPI (blue) of untransduced cells (no αSyn; top panel), and αSyn overexpressing cells (αSyn) without and with siRNA transfection. Scale bar: 50 µm. Panels on the right: illustration of the neuronal network as identified by the ‘neurite analyzer’ plugin in Fiji. The outline of the nuclei is highlighted in yellow. **e, f:** quantification of the total branch length (**e**) and the number of quadruple points as measure for network complexity (**f**) of the conditions illustrated in **d**. * *p* < 0.05, ** *p* < 0.01, *** *p* < 0.001, vs untransfected AV- αSyn cells, ^#^ *p* < 0.05, ^##^ *p* < 0.01, ^###^ *p* < 0.001 vs AV- αSyn cells transfected with a negative control siRNA. § *p* < 0.05, §§§ *p* < 0.001 vs. untransduced cells. **g**: Quantification of LDH, released into the cell culture medium as measure for cytotoxicity of control cells without αSyn overexpression (ctrl), untransfected αSyn overexpressing cells, and αSyn overexpressing cells transfected with siPOOL siRNA against SNX5 or negative control (neg ctrl) esiRNA.

### *SNX5* knockdown does not affect the expression of other retromer components

Since SNX5 protein is a part of the retromer complex (**Fig. 3a**) as part of SNX-BAR heterodimers build from SNX1/2 with SNX5/6, we investigated whether *SNX5* knockdown would lead to changes in the expression of other retromer components, as a possible compensatory mechanism. However, we did not observe alterations of VPS35, or one of the other components of the SNX-BAR heterodimers (SNX1, SNX2, SNX6) upon *SNX5* knockdown by a Western blot analysis (**Fig. 3b-f**). In addition, we performed knockdown experiments with siRNAs targeting these other components of the SNX-BAR-heterodimers and investigated whether the knockdown of these genes effects the cell viability. *SNX1* and *SNX2* knockdown did not alter the amount of released LDH, whereas *SNX6* knockdown resulted in an increased LDH release, indicating a toxic effect of the knockdown (**Fig. 3g**).

**Figure 3:**
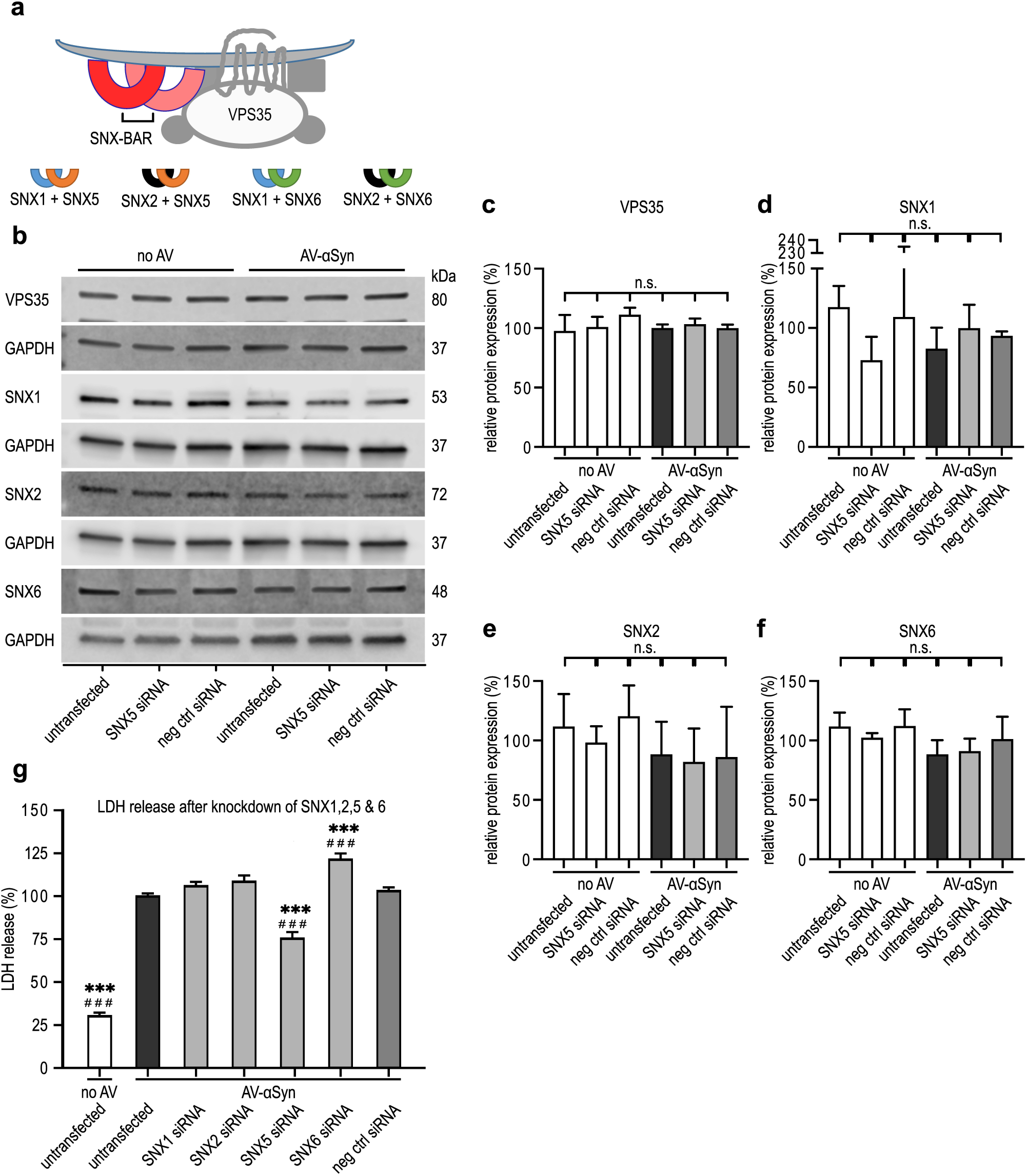
Investigation of the effect of the knockdown of other retromer components. **a**: Schematic illustration of the retromer complex. The SNX-BAR heterodimers are composed of SNX1 (blue arches) or SNX2 (black arches) in combination with SNX5 (orange arches) or SNX6 (green arches). **b**: Representative Western blots of other retromer components after staining with antibodies against VPS35, SNX1, SNX2, and SNX6. Below each blots the respective staining with an antibody against GAPDH used as loading control is shown. no AV: samples from untransduced cells, AV-αSyn: samples from cells with adenoviral mediated overexpression of αSyn. The cells were either untransfected or transfected with siRNA against SNX5 (SNX5 siRNA), or negative control siRNA (neg ctrl). **c-f**: Quantification of the Western blot bands. The white bars show data from untransduced cells, the grey bars αSyn overexpressing cells. Neither αSyn overexpression nor SNX5 knockdown (SNX5 siRNA) did significantly alter the protein expression levels of the retromer components VPS35, SNX1, SNX2, and SNX6. Data are shown as mean ± SEM. n.s.: not significant, one-way ANOVA with Tukey’s post-hoc test. **g**: Quantification of cytotoxicity in untransduced control cells (ctrl; white bar), in untransfected αSyn overexpressing cell and in αSyn overexpressing cells transfected with siRNAs against SNX1, SNX2, SNX5 and SNX6, or negative control siRNA. Knockdown of SNX1 and SNX2 had no significant effect on αSyn-induced toxicity. Knockdown of SNX5 protected against αSyn-induced toxicity. Knockdown of SNX6 increased αSyn-induced toxicity. Data are normalized to LDH release in untransfected αSyn overexpressing cells and shown as mean ± SEM. *** *p* < 0.0001 vs untransfected αSyn overexpressing cells (AV-αSyn). ### *p* < 0.0001 vs αSyn overexpressing cells transfected with a negative control siRNA.

### Knockdown of *SNX5* prevents the transport of αSyn into the trans-Golgi network

We next investigated the effect of *SNX5* knockdown on the transportation of αSyn within the cells (Fig. 4). Therefore, αSyn monomers were fluorescently labelled with ATTO-565 and added to the culture medium. The timeline is shown in **Fig. 4a**.

**Figure 4:**
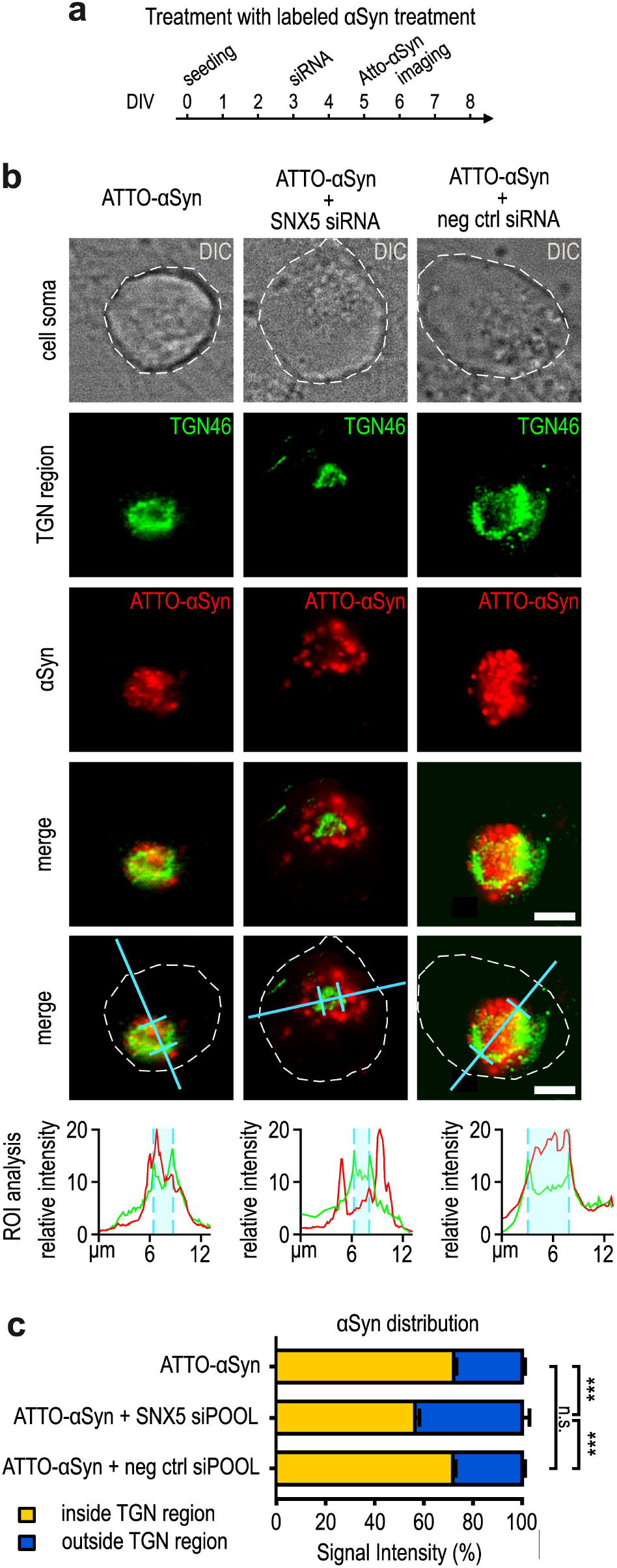
SNX5 knockdown prevents internalization and accumulation of exogenous αSyn into the trans-Golgi network. **a**: Experimental timeline of the treatment with fluorescently labelled αSyn via the culture medium. DIV: days in vitro. **b**: Immunofluorescence images of stainings with a TGN46 antibody (green) of cells treated with ATTO-565-labelled -αSyn (red) without knockdown of SNX5 (left side images), knockdown of SNX5 (images in the middle), and cells transfected with a negative control siRNA. The panel in the third row show merged images, the panels below show the same merged images with the hallmarks for quantification added. The turquoise lines indicate the location of the intensity measurement, the quantification of these measurements are shown in the graphs below. Lower panels: The turquoise area in the fluorescence intensity profiles indicate the outer margins of the TGN region, as defined by the TGN46 staining. The red line indicates the location of αSyn, both inside and outside of the TGN. After SNX5 knockdown, αSyn was distributed outside the TGN as indicated by the red intensity signal (images in the middle), which was not the case with the negative control siRNA (right side images). Scale bars 4 μm. **c**: Quantification of the proportion of the ATTO- αSyn inside and outside of the TGN region in the conditions shown in **b**. *** *p* < 0.001 ANOVA with Tukey’s post-hoc test. n.s. not significant.

After fixing the cells, we performed an immunohistrochemistry staining with a TGN antibody (TGN46), and performed a co-localization analysis between αSyn and the TGN area. The TGN area was defined from the profile of the fluorescence signal after TGN46 staining (**Fig. 4b, c**). We then quantified the fluorescence signal of ATT- αSyn inside and outside this region. By this, we could confirm that extracellularly added αSyn was indeed transported to the TGN region under normal conditions (**Fig. 4b**, left side images). Upon SNX5 knockdown, the proportion of ATTO-αSyn inside the TGN region compared to outside the TGN region was shifted towards less αSyn inside the TGN region (**Fig. 4b**, images in the middle). This effect that was not observed after transfection with a control siRNA (**Fig. 4b**, right side images). This suggests that SNX5 knockdown led to reduced trafficking of αSyn to the TGN.

### αSyn leads to TGN fragmentation which is ameliorated by *SNX5* knockdown

The morphology of the TGN can be divided in three states (normal, scattered, fragmented) (17). The different TGN morphologies observed in LUHMES cells exposed to extracellular ATTO-αSyn are shown in **Fig. 5a**, their relative quantitative frequency in **Fig. 5e**. The TGN morphologies in cells transduced to overexpress αSyn (AV-αSyn) are illustrated in **Fig. 5b** with quantitative data in **Fig. 5e**.

**Figure 5:**
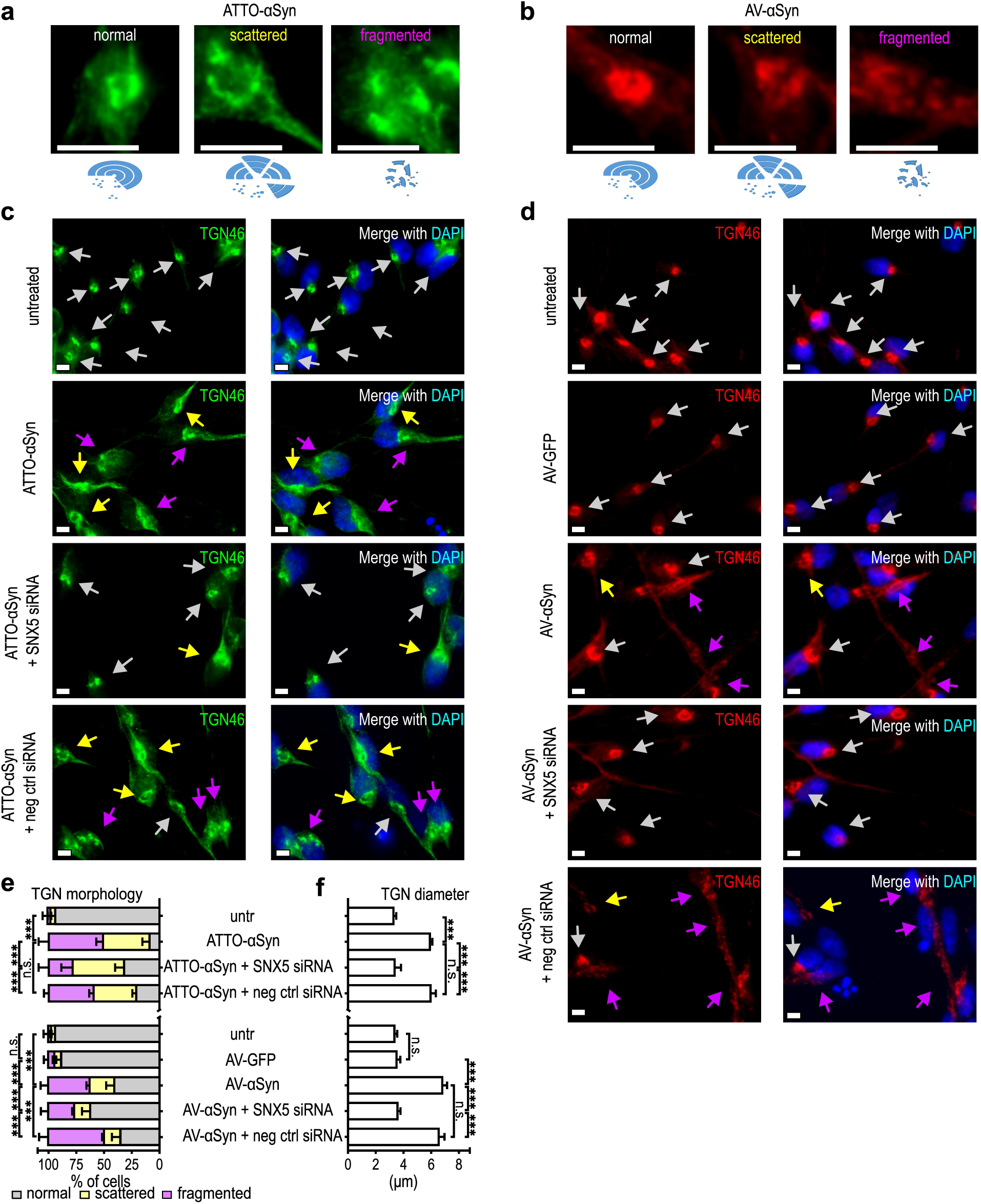
SNX5 knockdown prevents TGN scattering and fragmentation in LUHMES cells. **a**: Images of LUHMES cells treated with ATTO-565-labeled recombinant αSyn monomers, stained with an antibody against TGN46 (green), to illustrate different states of TGN morphology. The left, middle and right panels show normal, scattered, and fragmented TGN morphology, respectively (schematically illustrated in the sketches below the respective panels). White arrows: normal morphology, yellow arrows: scattered, purple arrows: fragmented. All these morphologies were observed in αSyn-treated cells, but to varying degrees. **b**: Images of LUHMES cells transduced with αSyn-overexpressing adenoviral vectors (AV), stained with an antibody against TGN46 (red) to illustrate the same states of TGN morphology as in **a**. **c**: Representative images of stainings with an antibody against TGN46 (green; left), and merged with a DAPI staining (blue; right), of untreated cells, untransfected ATTO-αSyn-treated cells, ATTO-αSyn-treated cells with SNX5 siRNA transfection, and ATTO-αSyn-treated cells with negative control (neg ctrl) siRNA transfection. The arrows indicate the different states of TGN morphology as illustrated in **a**. **d**: Representative images of LUHMES cells stained with an antibody against TGN46 (red; left) and merged with a DAPI staining (blue; right) of untransduced/untransfected control cells (ctrl), untransfected cells transduced with an adenovirus to overexpress GFP (AV-GFP) or to overexpress αSyn (AV- αSyn), and αSyn-overexpressing cells with SNX5 siRNA or negative control (neg ctrl) siRNA transfection. **e**: Percentage of cells with normal (grey), scattered (yellow), or fragmented (purple) TGN morphology in the experimental conditions of **c** and **d**. Exogenous αSyn (left) as well as adenoviral overexpressed αSyn (right) led to a higher percentage of scattered or fragmented TGNs, which was ameliorated by SNX5 knockdown. **f**: Quantification of the TGN diameter. Both, exogenous αSyn (ATTO- αSyn) and adenoviral overexpressed αSyn (AV-αSyn) led to larger TGN sizes. *** *p* < 0.001. ANOVA with Tukey’s post hoc test. n.s. not significant.

Then, we studied the effect of *SNX5* knockdown on this effect induced by ATTO-αSyn (**Fig. 5c**) and AV-αSyn, using AG-overexpression of GFP as control protein (**Fig. 5d**).

The quantification of these experiments demonstrated that both exposure to extracellular ATTO-αSyn and intracellular AV-αSyn-overexpression, but not AV-GFP-overexpression, led to a significant increase in scattered and fragmented TGN, as compared to untreated control cells, and that this effect was partially prevented by SNX5-knockdown, but not by a control siRNA (**Fig. 5e**).

Furthermore, we observed that ATTO-αSyn treatment led to an increase of the TGN diameter from 3.4 ± 0.1 μm (untreated controls) to 6.1 ± 0.2 μm (p<0.001). Overexpression of αSyn led to an increase to 6.8 ± 0.2 μm (p<0.001). However, the knockdown of SNX5 led to a prevention of this increase of the TGN diameter in ATTO-αSyn treated cells to 3.5 ± 0.2 μm and in AV- αSyn transduced cells to 3.6 ± 0.1 μm, with no more significant difference to untreated or AV- GFP transduced cells (3.6 ± 0.1 µm; **Fig. 5f**).

### *SNX5* knockdown led to increased levels of αSyn in early endosome

To further investigate the mechanism between *SNX5* knockdown and protection from αSyn-induced toxicity, we examined the endosome to TGN pathway, the endosome to plasma membrane (PM; recycling endosome) pathway, the endosome to lysosome pathway, and the autophagosome to lysosome pathway.

Therefore, LUHMES cells were exposed to extracellular ATTO-αSyn either with or without *SNX5* knockdown, prior to fixation for immunocytochemistry. We used an antibody against Rab5a as a marker for the early endosome, antibodies against Rab7 and LAMP1 as markers for the late endosome, an antibody against LAMP2a as marker for the lysosomes, antibodies against p62 and LC3B as markers for autophagosomes, and an antibody against Rab11a as marker for recycling endosomes (**Fig. 6a**). Manderson’s overlap coefficient was determined to evaluate the degree of co-localization between αSyn and the different vesicular structures (**Fig. 6b**).

**Figure 6:**
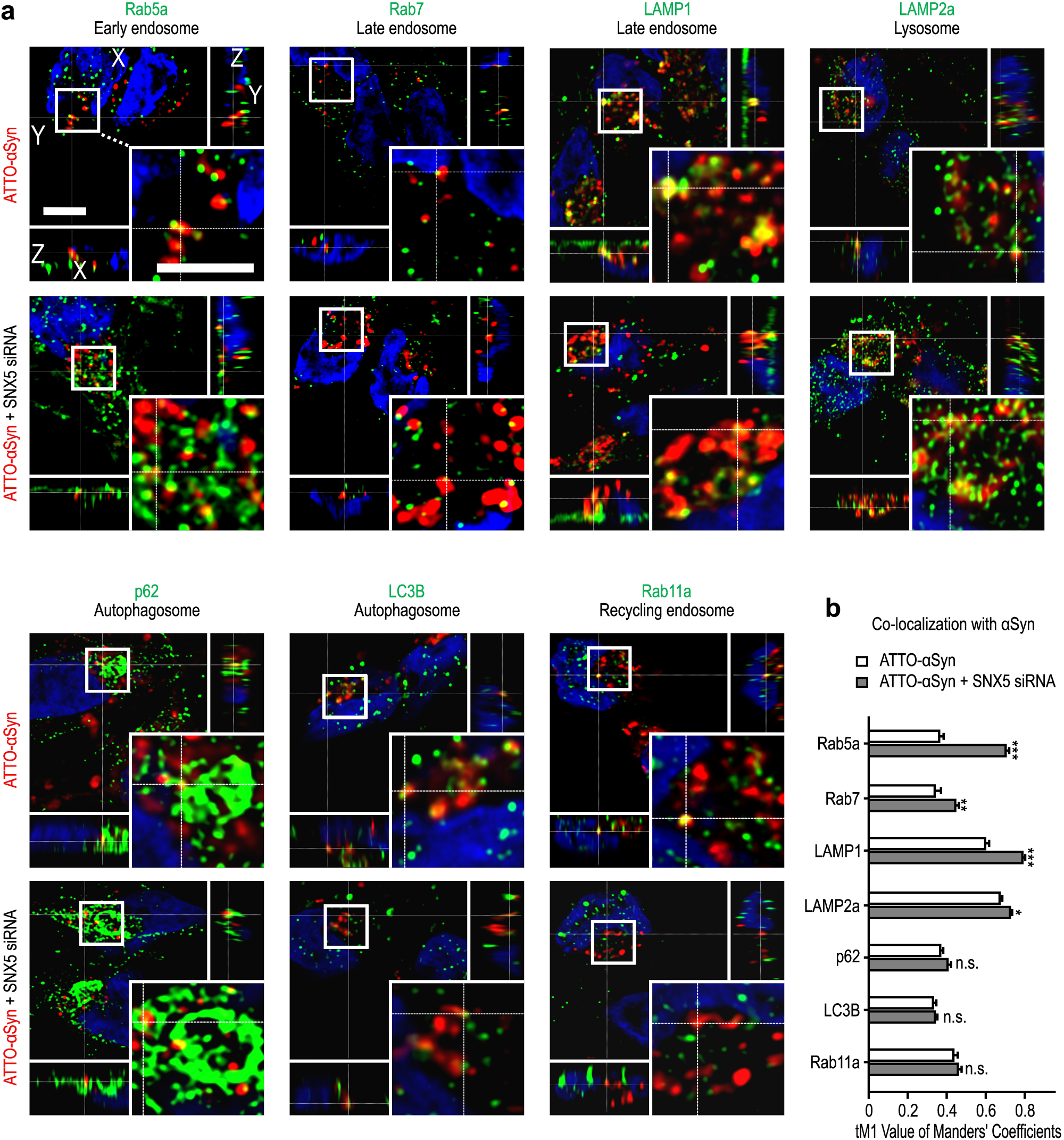
Co-localization of internalized αSyn with endocytosis markers. **a:** Representative images of LUHMES cells treated with fluorescently labeled αSyn (ATTO- αSyn, red) with control siRNA transfection (left column of images), and after knockdown of either SNX5 (middle column of images) or SNX6 (right column of images). The markers used in this experiment are indicated in green left of each the panel. A selected region of each image (white square) is shown in higher magnification in the right bottom corner. Scale bars: 4 µm. **b:** Quantification of the degree of co-localization between exogenous αSyn and endocytosis markers. SNX5 knockdown (light grey bars) led to increased co-localization between αSyn and early endosomes (Rba5a), late endosomes (Rab7, LAMP1), and lysosomes (LAMP2A) compared to untransfected cells (white bars). SNX6 knockdown on the other hand led to a slight reduction of co-location between αSyn and early endosomes (Rab5a) and recycling endosomes (Rab11a). Data are represented as mean ± SEM. * *p* < 0.05, ** *p* < 0.01, *** *p* < 0.001 vs ATTO- αSyn, ^#^ *p* < 0.05, ^##^ *p* < 0.01, ^###^ *p* < 0.001 vs ATTO- αSyn + SNX5 siRNA; n.s. not significant; one-way ANOVA with Tukey’s post hoc test.

Under normal conditions, αSyn was more co-localized (Manderson’s overlap coefficient > 0.5) with endosomes and lysosome than with autophagosomes or the recycling endosomes, suggesting that αSyn was mainly transported via the endosome to lysosome pathway and accumulated later in lysosomes under normal conditions.

*SNX5* knockdown led to a strong increase of co-localization between αSyn and early endosomes (Rab5), from 0.4 ± 0.02 to 0.77 ± 0.02 (p<0.001) and to a slight increase of the co-localization between αSyn and the late endosome markersRab7: (from 0.37 ± 0.04 to 0.49 ± 0.02; p<0.001) and LAMP1 (from 0.66 ± 0.02 to 0.86 ±0.02; p<0.001). On the other hand, co-localization to autophagosomes and lysosomes was not altered (**Fig. 6b**). Together with the observation that *SNX5* knockdown led to reduced abundance of αSyn in the TGN (**Fig. 4**), these observations suggest that SNX5 facilitates αSyn-trafficking from the early endosome to the TGN (**Fig. 7a**) and that *SNX5* knockdown reduces this transport and leads to more αSyn in early and late endosomes (**Fig 7b**).

**Figure 7.**
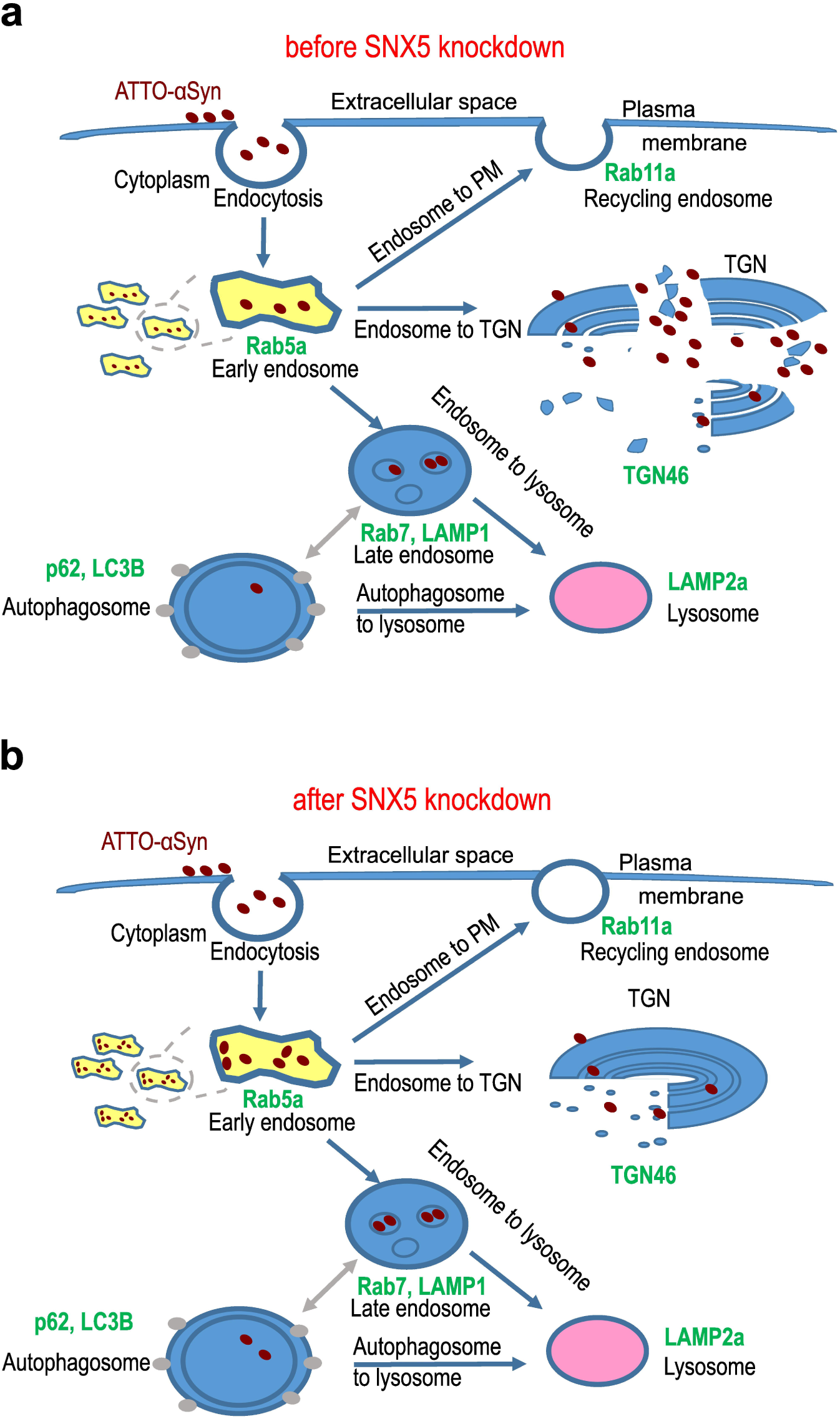
Transportation of αSyn before and after knockdown of SNX5. **a, b:** Schematic illustration of possible endocytic pathways and the markers of the different compartment used in our study (green). After endocytosis from the extracellular space, αSyn (red dots) is transported are transported into the early endosomes (yellow; Rab5a). These are then either transported back to the plasma membrane (endosome to PM) in form of recycling endosomes (Rab11), to the trans-Golgi network (TGN; TGN46; endosome to TGN) or form late endosomes (Rab7, LAMP1), which are then either transported directly to lysosomes (LAMP2a; endosome to lysosome) or first to autophagosomes (p62, LC3B) and then to lysosomes (autophagosome to lysosome). Before SNX5 knockdown, more αSyn is transported to the TGN (**c**), while after SNX5 knockdown less αSyn is transported to the TGN, but more the early and late endosomes (**d**).

## Discussion

In the present work, we conducted a genome-wide multi-step RNAi screening in a PD human neuronal cell model, in order to identify genes, whose knockdown protected from αSyn-induced toxicity. From 69 hits that were identified in the primary screening, 28 were confirmed in the secondary screening. After exclusion of genes, the knockdown of which unspecifically protected the cells as determined by their effect in GFP expressing control cells, 12 genes remained, whose knockdown specifically protected from alpha-synuclein induced toxicity, with 4 remaining significant after correction for multiple testing. Amongst these, sorting nexin 5 (SNX5), a component of the retromer complex, was the most promising candidate to follow-up. We first confirmed the protective efficacy of SNX5 knockdown with a second RNAi system. Since SNX5 is part of the SNX-BAR heterodimer within the retromer, we investigated, if SNX5 knockdown led to the compensatory regulation of other retromer components, which was not the case. However, we found that in contrast to SNX5 knockdown, the knockdown of an SNX6, the alternative for SNX5 in a possible composition of SNX-BAR heterodimers increased αSyn-induced toxicity, suggesting that SNX5 was involved in trafficking of toxic αSyn species. We found that αSyn overexpression and treatment of the cells with exogenous αSyn led to fragmentation of the trans-Golgi network (TGN) and that SNX5 knockdown prevented TGN fragmentation, by inhibiting the transport of αSyn to the TGN. Our findings suggest that SNX5 is involved in the regulation of the trafficking and toxicity of αSyn and could be a promising target for the future development of neuroprotective therapies for PD and related synucleinopathies.

High-throughput RNA interference (RNAi) screenings are an effective way to identify new genes that are involved in the pathophysiology of distinct conditions in a hypothesis-free way. However, in the past RNAi screenings that investigated modifiers of αSyn toxicity were performed only in subsets of the genome and/or in non-human cells (for review see: (18)). To our knowledge, we present the first whole-genome RNAi screening of modifier αSyn-induced toxicity. Furthermore, we utilized human postmitotic dopaminergic neurons (LUHMES cells), which very closely resemble the cells, the demise of which is responsible for the motor symptoms of PD patients. For the screening, we used a multistep approach to ensure that our final hits specifically modified αSyn-induced toxicity.

On the one hand, SNX5 was the hit with the lowest p-value in the confirmatory screening process. On the other hand, as part of the SNX-BAR heterodimer which itself is part of the retromer, SNX5 was particularly interesting. It is known that the retromer plays a role in the pathophysiology of PD (19–21). Mutations in an important component of the retromer, VPS35, are risk factors to develop PD, but the frequency of such mutations is not yet known (22, 23). Moreover, the retromer is involved in multiple cellular processes including vesicular trafficking, receptor recycling, but also mitochondrial function and dopamine signaling (22). In addition, it is currently not yet fully understood, if VPS35 mutation lead to a loss or gain of function of the retromer (23). Another part of the retromer complex are SNX-BAR heterodimers (see Figure 3A). These are composed of SNX1 or SNX2 in combination with SNX5 and SNX6 (24). The composition of the SNX-BAR heterodimers plays a role in cargo-sorting (25).

Therefore, we hypothesized that SNX5 was involved in trafficking of toxic αSyn species. This hypothesis was supported by the fact that knockdown of SNX6, the alternative to SNX5 in the SNX-BAR heterodimers increased toxicity, whereas knockdown of other retromer components had no influence on cell viability.

We then investigated, the intracellular localization of exogenously added αSyn in our model and found that upon uptake αSyn was co-localized with the TGN, suggesting a transportation of αSyn to the TGN in our cell model. In line with that also others showed that αSyn co-localizes with the TGN in human astrocytes (26) and in neuroblastoma cells (27). Furthermore, we observed fragmentation of the TGN as consequence of αSyn overexpression and treatment with exogenous αSyn. Consistently, others found that treatment with prefibrillar αSyn aggregates led to TGN fragmentation in an immortalized fibroblast cell line derived from monkey kidneys (28). Furthermore, fragmentation of the TGN has been previously described in nigral neurons of PD patients (29). Moreover, also in other PD cell models, αSyn overexpression led to Golgi fragmentation (30, 31).

Together these findings emphasize that our cell model reflects what is observed in patients and supports the relevance of our findings in the present study. Interestingly, SNX5 knockdown prevented trafficking of αSyn to the TGN and its fragmentation, suggesting that SNX5 is a key regulator in trafficking of toxic αSyn species. In the past, also others could show that trafficking of αSyn plays an important role. Previously, in a yeast model it was shown that αSyn blocked the ER-Golgi transport. The authors showed that overexpression of the orthologue of Rab1, a protein needed for the docking of transport vesicles with the Golgi apparatus and thus improving the TGN function protected dopaminergic neurons from αSyn induced toxicity in yeast (32). In line with this study, we could also show that impairment of the TGN is toxic for dopaminergic neurons.

While restoration of the ER-Golgi traffic is one possibility to prevent toxicity, our data suggest that also redirection of harmful αSyn species away from the TGN by knockdown of *SNX5* could be a strategy to prevent αSyn-included cell death. Interestingly, we could previously show that αSyn was degraded by macroautophagy and that stimulation of autophagy (8) but that also bypassing macroautophagy by prevention of the formation of autophagosomes protected from αSyn-induced toxicity (7), emphasizing that in some situations the stimulation and bypassing of a distinct intracellular pathway can both be protective. Furthermore, others have also demonstrated the importance of αSyn. In H4 neuroglial cells it was shown that the knockdown of distinct Rab GTPases that were involved in αSyn trafficking prevented the formation of αSyn inclusions (33).

## Conclusion

In summary, we performed a genome-wide siRNA screening and identified *SNX5* as the top hit. We found that SNX5 protein as part of the retromer complex was involved in αSyn trafficking. We found that αSyn was taken up and transported to the TGN and led to its fragmentation. Both could be ameliorated by knockdown of *SNX5*. On the other hand, SNX6 knockdown was toxic for the cells, suggesting that a distinct regulation of intracellular αSyn trafficking involves SNX-BAR heterodimers. Furthermore, our study emphasizes that further investigation in αSyn trafficking could lead to the development of new therapeutic options to save neurons from αSyn-induced cell death. In particular, SNX5 seems to be a promising novel target for the development of a neuroprotective treatment for PD and related synucleinopathies.

## List of abbreviations

aa: Amino Acids
αSyn: Alpha-Synuclein
ATTO-αSyn: ATTO-labelled αSyn
AV-αSyn: overexpressing αSyn
bFGF: Basic Fibroblast Growth Factor
BAR: Bin/Amphiphysin/Rvs
DAPI: 4′,6-diamidino-2-phenylindole
DFGF: Basic Fibroblast Growth Factor
DLB: Dementia with Lewy Bodies
DMEM/F12: Dulbecco’s Modified Eagle Medium/Nutrient Mixture F-12
DM: Differentiation Medium
ECL: Enhanced Chemiluminescence
esiRNAs: endoribonuclease-prepared small interfering RNAs
GAPDH: Glyceraldehyde 3-phosphate Dehydrogenase
GCI: Glial Cytoplasmic Inclusions
GFP: Green Fluorescent Protein
GM: Growth Medium
GDNF: Glial cell-derived Neurotrophic Factor
HBSS: Hanks’ Balanced Salt Solution
HRP: Horse Radish Peroxidase
ICC: immunocytochemistry
LDH: Lactate Dehydrogenase
LUHMES: Lund Human Mesencephalic Cells
MOI: Multiplicity of Infection
N2: Supplement N2
NADH: Nicotinamide Adenine Dinucleotide
NHS: Normal Horse Serum
PBS: Phosphate Buffered Saline
PD: Parkinson’s Disease
PFA: Paraformaldehyde
PM: Plasma Membrane
PVDF: Polyvinylidene Difluoride
PI: Propidium Iodide
qPCR: Quantitative Polymerase Chain Reaction
RNAi: High-throughput RNA interference
ROI: Region of Interest
SDS-PAGE: Sodium Dodecyl Sulfate Polyacrylamide Gel Electrophoresis
SNX5: Sorting Nexin 5
TBS-T: Tris-buffered Saline with Tween-20
TGN: Trans Golgi Network
VPS35: Vacuolar Protein Sorting Ortholog 35

## Declaration

### Ethics approval and consent to participate

N/A

### Consent for publication

N/A

### Availability of data and materials

All datasets used and analyzed in this study are available from the corresponding authors on reasonable request.

### Authors’ contributions

MH, LD, OWC, CM, SB performed the experiments, FH and CWS reviewed the experimental design, MH, MC, and GUH conceived the study, MH and LD created the figures. MH wrote the first draft of the manuscript. All authors were involved in editing, correction, and finalizing the manuscript

## Acknowledgements

We thank Sabine Lang (MHH, Dept. for Neurology) for technical support.

## Literature Cited

1. Goedert M, Spillantini MG, Del Tredici K, Braak H. 100 years of Lewy pathology. Nat Rev Neurol 2013; 9(1):13–24.

2. Fanciulli A, Stankovic I, Krismer F, Seppi K, Levin J, Wenning GK. Multiple system atrophy. Int Rev Neurobiol 2019; 149:137–92.

3. Tofaris GK. Initiation and progression of α-synuclein pathology in Parkinson’s disease. Cell Mol Life Sci 2022; 79(4):210.

4. Burré J, Sharma M, Südhof TC. Cell Biology and Pathophysiology of α-Synuclein. Cold Spring Harb Perspect Med 2018; 8(3).

5. Lotharius J, Falsig J, van Beek J, Payne S, Dringen R, Brundin P et al. Progressive degeneration of human mesencephalic neuron-derived cells triggered by dopamine-dependent oxidative stress is dependent on the mixed-lineage kinase pathway. Journal of Neuroscience 2005; 25(27):6329–42.

6. Höllerhage M, Goebel JN, Andrade A de, Hildebrandt T, Dolga A, Culmsee C, et al. Trifluoperazine rescues human dopaminergic cells from wild-type α-synuclein-induced toxicity. Neurobiol Aging 2014; 35(7):1700–11.

7. Fussi N, Höllerhage M, Chakroun T, Nykänen N-P, Rösler TW, Koeglsperger T et al. Exosomal secretion of α-synuclein as protective mechanism after upstream blockage of macroautophagy. Cell Death and Disease 2018; 9(7):757.

8. Höllerhage M, Fussi N, Rösler TW, Wurst W, Behrends C, Höglinger GU. Multiple molecular pathways stimulating macroautophagy protect from alpha-synuclein-induced toxicity in human neurons. Neuropharmacology 2019; 149:13–26.

9. Höllerhage M, Moebius C, Melms J, Chiu W-H, Goebel JN, Chakroun T et al. Protective efficacy of phosphodiesterase-1 inhibition against alpha-synuclein toxicity revealed by compound screening in LUHMES cells. Sci Rep 2017; 7(1):11469.

10. Harbour ME, Breusegem SYA, Antrobus R, Freeman C, Reid E, Seaman MNJ. The cargo-selective retromer complex is a recruiting hub for protein complexes that regulate endosomal tubule dynamics. Journal of Cell Science 2010; 123(Pt 21):3703–17.

11. Rodriguez L, Marano MM, Tandon A. Import and Export of Misfolded α-Synuclein. Front Neurosci 2018; 12:344.

12. Follett J, Norwood SJ, Hamilton NA, Mohan M, Kovtun O, Tay S et al. The Vps35 D620N mutation linked to Parkinson’s disease disrupts the cargo sorting function of retromer. Traffic 2014; 15(2):230–44.

13. Kittler R, Surendranath V, Heninger A-K, Slabicki M, Theis M, Putz G et al. Genome-wide resources of endoribonuclease-prepared short interfering RNAs for specific loss-of-function studies. Nat Methods 2007; 4(4):337–44.

14. Theis M, Buchholz F. MISSION esiRNA for RNAi screening in mammalian cells. J Vis Exp 2010; (39).

15. Haas AJ, Prigent S, Dutertre S, Le Dréan Y, Le Page Y. Neurite analyzer: An original Fiji plugin for quantification of neuritogenesis in two-dimensional images. J Neurosci Methods 2016; 271:86–91.

16. Hoffmann A-C, Minakaki G, Menges S, Salvi R, Savitskiy S, Kazman A et al. Extracellular aggregated alpha synuclein primarily triggers lysosomal dysfunction in neural cells prevented by trehalose. Sci Rep 2019; 9(1):544.

17. Paiva I, Jain G, Lázaro DF, Jerčić KG, Hentrich T, Kerimoglu C et al. Alpha-synuclein deregulates the expression of COL4A2 and impairs ER-Golgi function. Neurobiology of disease 2018; 119:121–35.

18. Höllerhage M, Bickle M, Höglinger GU. Unbiased Screens for Modifiers of Alpha-Synuclein Toxicity. Curr Neurol Neurosci Rep 2019; 19(2):8.

19. Cui Y, Yang Z, Teasdale RD. The functional roles of retromer in Parkinson’s disease. FEBS Lett 2018; 592(7):1096–112.

20. Yang Z, Li Z, Teasdale RD. Retromer dependent changes in cellular homeostasis and Parkinson’s disease. Essays Biochem 2021; 65(7):987–98.

21. Williams ET, Chen X, Moore DJ. VPS35, the Retromer Complex and Parkinson’s Disease. J Parkinsons Dis 2017; 7(2):219–33.

22. Rahman AA, Morrison BE. Contributions of VPS35 Mutations to Parkinson’s Disease. Neuroscience 2019; 401:1–10.

23. Sassone J, Reale C, Dati G, Regoni M, Pellecchia MT, Garavaglia B. The Role of VPS35 in the Pathobiology of Parkinson’s Disease. Cell Mol Neurobiol 2021; 41(2):199–227.

24. Simonetti B, Danson CM, Heesom KJ, Cullen PJ. Sequence-dependent cargo recognition by SNX-BARs mediates retromer-independent transport of CI-MPR. J Cell Biol 2017; 216(11):3695–712.

25. Kvainickas A, Jimenez-Orgaz A, Nägele H, Hu Z, Dengjel J, Steinberg F. Cargo-selective SNX-BAR proteins mediate retromer trimer independent retrograde transport. J Cell Biol 2017; 216(11):3677–93.

26. Rostami J, Holmqvist S, Lindström V, Sigvardson J, Westermark GT, Ingelsson M et al. Human Astrocytes Transfer Aggregated Alpha-Synuclein via Tunneling Nanotubes. Journal of Neuroscience 2017; 37(49):11835–53.

27. Hivare P, Gadhavi J, Bhatia D, Gupta S. α-Synuclein fibrils explore actin-mediated macropinocytosis for cellular entry into model neuroblastoma neurons. Traffic 2022; 23(7):391–410.

28. Gosavi N, Lee H-J, Lee JS, Patel S, Lee S-J. Golgi fragmentation occurs in the cells with prefibrillar alpha-synuclein aggregates and precedes the formation of fibrillar inclusion. Journal of Biological Chemistry 2002; 277(50):48984–92.

29. Fujita Y, Ohama E, Takatama M, Al-Sarraj S, Okamoto K. Fragmentation of Golgi apparatus of nigral neurons with alpha-synuclein-positive inclusions in patients with Parkinson’s disease. Acta Neuropathologica 2006; 112(3):261–5.

30. Martínez-Menárguez JÁ, Tomás M, Martínez-Martínez N, Martínez-Alonso E. Golgi Fragmentation in Neurodegenerative Diseases: Is There a Common Cause? Cells 2019; 8(7).

31. Nakagomi S, Barsoum MJ, Bossy-Wetzel E, Sütterlin C, Malhotra V, Lipton SA. A Golgi fragmentation pathway in neurodegeneration. Neurobiology of disease 2008; 29(2):221–31.

32. Cooper AA, Gitler AD, Cashikar A, Haynes CM, Hill KJ, Bhullar B et al. Alpha-synuclein blocks ER-Golgi traffic and Rab1 rescues neuron loss in Parkinson’s models. Science 2006; 313(5785):324–8.

33. Gonçalves SA, Macedo D, Raquel H, Simões PD, Giorgini F, Ramalho JS et al. shRNA-Based Screen Identifies Endocytic Recycling Pathway Components That Act as Genetic Modifiers of Alpha-Synuclein Aggregation, Secretion and Toxicity. PLoS Genet 2016; 12(4):e1005995.

